# Ori-Finder-Arch: An Updated Web Server for the Annotation and Visualization of Archaeal Replication Origins

**DOI:** 10.64898/2026.08.15.744077

**Authors:** Zhisong You, Zetong Zhang, Hao Luo, Feng Gao

**Affiliations:** Department of Physics, School of Science, Tianjin University, Tianjin, 300072, China; State Key Laboratory of Synthetic Biology, Tianjin University, Tianjin, 300072, China; Frontiers Science Center for Synthetic Biology and Key Laboratory of Systems, Bioengineering (Ministry of Education), Tianjin University, Tianjin, 300072, China

**Keywords:** Archaea, DNA replication origin, Replication initiation protein, Z-curve method, Web server

## Abstract

Archaea are promising chassis organisms in biotechnology, and the accurate annotation of their chromosomal replication origins (*oriC*s) is the key to unlocking their full potential. However, the existing Ori-Finder 2 web server suffers from low accuracy, slow speed, and limited scalability. In this study, we present Ori-Finder-Arch, an updated web server for high-performance *oriC* prediction in archaea. This pipeline integrates HMMER-based replication initiation protein (RIP) annotation, refined consensus motif recognition, and GC profile-based DNA unwinding element (DUE) detection. On a benchmark set of experimentally validated *oriC*s, Ori-Finder-Arch achieved a recall of 95.6% and a precision of 86.0%, substantially outperforming Ori-Finder 2 (62.2% and 63.6%, respectively), while running 4.75 times faster and supporting diverse assembly levels. When applied to the available archaeal assemblies, it successfully annotated 17,472 *oriC*s. Meanwhile, the web server provides interactive visualizations at different levels. In conclusion, Ori-Finder-Arch offers an efficient, accurate, and user-friendly platform for advanced studies of archaeal DNA replication initiation and synthetic biology applications, and is freely available at https://tubic.org/Ori-Finder-Arch/ and https://tubic.tju.edu.cn/Ori-Finder-Arch/.

## Introduction

As a unique branch of the evolutionary tree, archaea often exhibit remarkable adaptability to various extreme environments (e.g., high temperature, low pH, and high salinity) and unique metabolic pathways that are not found in other organisms. These traits render archaea promising chassis organisms in biotechnology [1], and studying the DNA replication process in archaea, particularly the mechanism of replication initiation, is important for unlocking the full potential of the archaeal chassis. In prokaryotes, the initiation of DNA replication occurs at specific genomic loci known as chromosomal replication origins (*oriC*s) [2,3]. In bacteria, the chromosome typically contains a single *oriC*, whereas in archaea, the number of *oriC*s varies substantially among different species [4].

The experimental methods for identifying replication origins include neutral/neutral two- dimensional agarose gel electrophoresis (2D-AGE) [5], microarray-based marker frequency analysis (MFA) [6], and autonomously replicating sequence (ARS) assays [7]. These methods are often complemented by auxiliary techniques, such as chromatin immunoprecipitation (ChIP) [8] and electrophoretic mobility shift assay (EMSA) [9]. These techniques have facilitated the identification of multiple *oriC*s in archaea, including *H. volcanii* [10] and *S. islandicus* [11]. However, the development of metagenomic sequencing has led to an exponential expansion of genome sequences [12], exceeding the capacity of experimental methods.

Structurally, archaeal *oriC*s are typically located in intergenic regions (IGRs) adjacent to genes encoding replication initiation proteins (RIPs) and generally contain conserved consensus motifs and AT-rich regions [13] (Fig. 1). These motifs serve as origin recognition boxes (ORBs)- specific binding sites for RIPs, whereas the AT-rich regions serve as DNA unwinding elements (DUEs) [14]. For example, in *Sulfolobus*, a pair of inverted ORBs with a DUE located between them is required to recruit a pre-replication complex (pre-RC) and unwind DNA [15]. These features provide a biological foundation for the computational annotation of archaeal *oriC*s.

**Fig. 1.**
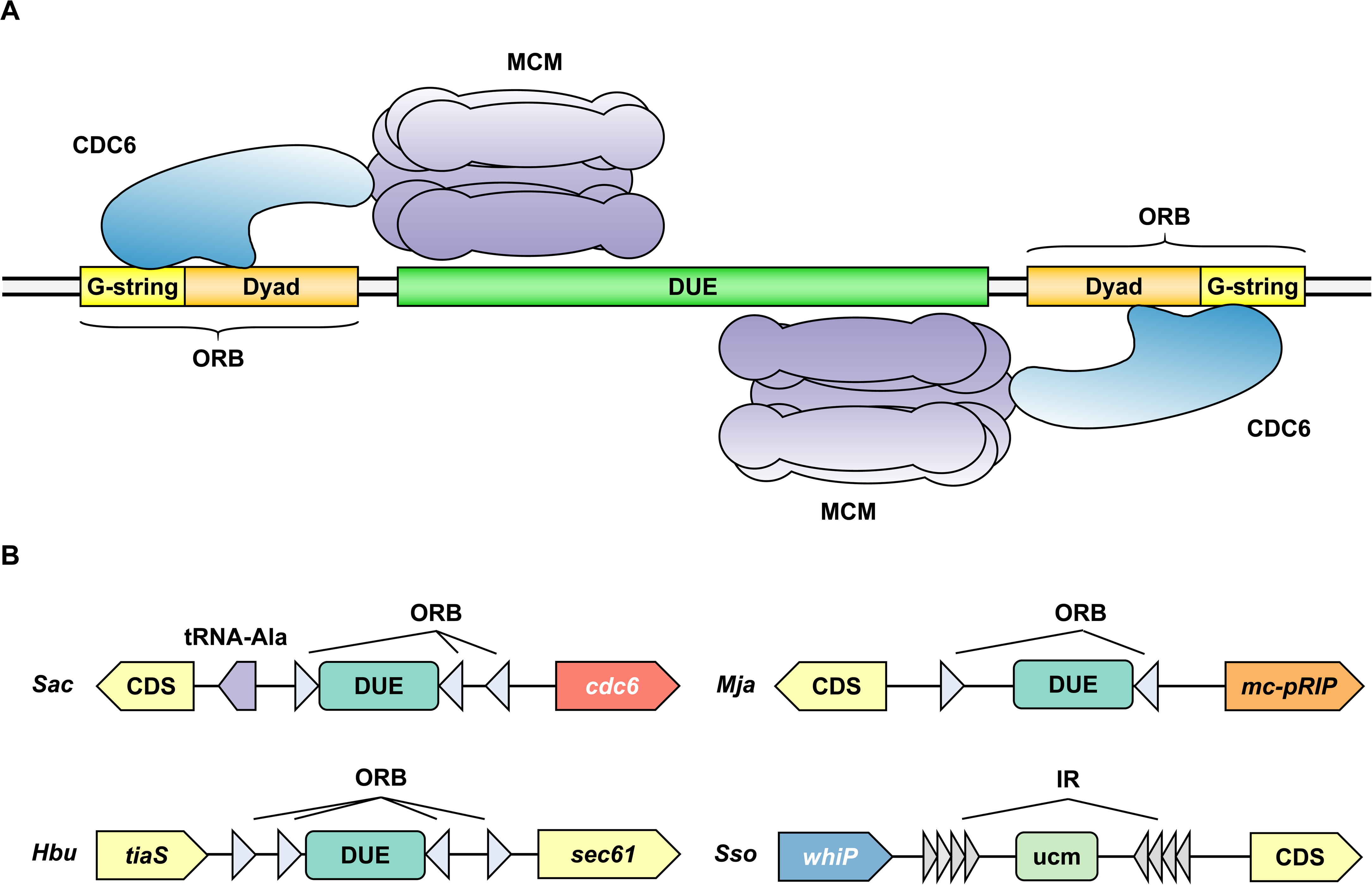
Typical structures of archaeal replication origins. **A.** The minimal structure required for the physiological function of an archaeal *oriC* (ORB(−)/DUE/ORB(+) pattern). During replication initiation, a pair of inverted ORBs recruit CDC6, which subsequently forms a complex with MCM to unwind the DNA in the intervening DUE region, thereby launching bidirectional replication. **B.** The typical *oriC* structures of four model organisms. The main *oriC*s of *S. acidocaldarius* (*Sac*), *M. jannaschii* (*Mja*), and *H. butylicus* (*Hbu*) all conform to the ORB(−)/DUE/ORB(+) pattern, and the only difference lies in the adjacent gene (note that *Hbu oriC* has no adjacent RIP gene). In contrast, the pattern of WhiP-type *oriC* is considerably different. The WhiP-type *oriC* of *S. solfataricus* (*Sso*) adopts an IR(+)/ucm/IR(−) pattern, implying a replication initiation mechanism entirely distinct from the former.

In our previous study, Ori-Finder 2, the only available web server for the computational annotation of archaeal *oriC*s [16], was developed. By integrating ZCURVE [17] and Glimmer3 [18] for gene recognition, BLAST+ [19] for RIP recognition, FIMO [20] for ORB identification, and the Z-curve method for visualization [21], Ori-Finder 2 has been used to annotate *oriC*s in newly sequenced archaeal genomes [22−26]. For example, four of six putative *oriC*s predicted by Ori-Finder 2 in *Natrinema* sp. J7 were experimentally verified [27]. Furthermore, Ori-Finder 2 has facilitated discoveries of shared evolutionary transitions and adaptations between marine bacteria and archaeal organisms [28], as well as the unique starvation survival systems in *Methanothermobacter thermautotrophicus* ΔH [29]. However, Ori-Finder 2 has several limitations. First, its accuracy remained low, as benchmarking against the DoriC 5.0 [30] database yielded a sensitivity of 66.7% and precision of 62.1%. Second, the server suffers from a slow processing speed and lacks a standalone version, making it unsuitable for large-scale processing. Third, the input and output options are restricted (e.g., genome assemblies with multiple sequences cannot be properly processed and no result download is supported), and only complete genomes are accepted.

Therefore, we introduced Ori-Finder-Arch, an updated web server for archaeal *oriC* annotation and visualization. By thoroughly investigating the structural features of *oriC*s and their associated elements and integrating state-of-the-art bioinformatics tools, we developed an updated, efficient, and accurate annotation pipeline for archaeal *oriC*s. On the benchmark dataset, Ori-Finder-Arch significantly outperformed Ori-Finder 2 in terms of running speed, recall, and precision. Moreover, Ori-Finder-Arch is equipped with an improved interactive visualization interface and enhanced input/output support, resulting in better usability and user-friendliness. We believe that this work provides a robust platform for the high-throughput prediction of *oriC*s, thereby advancing the mechanistic understanding of archaeal DNA replication initiation and supporting the rational design of archaeal chassis in synthetic biology.

## Methods

### Data Collection

#### Genomes

On January 8, 2026, 16,409 archaeal genomes spanning four assembly levels were downloaded from the NCBI Genome database [31], including 918 complete/chromosome-level and 15,491 scaffold/contig-level assemblies (Table S1). For each genome, plasmid sequences were removed using in-house Python scripts, and the taxonomic lineage was standardized using TaxonKit v0.20.0 [32]. Additionally, to assess the impact of assembly continuity on prediction, the scaffold/contig-level assemblies were stratified by N50 into three ranks here: high (N50 > 50 kb), medium (10 kb ≤ N50 ≤ 50 kb), and low (N50 < 10 kb).

#### *oriC*s and RIPs

Experimentally verified *oriC*s were compiled from the literature, yielding 45 items from 19 strains, which served as the benchmark dataset (Table S2). Additional *oriC*s sequences were retrieved from the DoriC 12.0 database [33], comprising 1,124 sequences from 662 strains. After removing entries that overlapped with the benchmark dataset, the remaining 1,080 sequences were designated as the analysis dataset. RIP sequences were initially derived from DoriC and then expanded against UniProtKB [34] using CD-search [35] and BLASTp [19]. After quality control, 864 reference sequences were retained (Table S3).

### Replication Initiation Protein Annotation

RIPs in archaea exhibit considerable functional domain diversity, and their accurate annotation requires tailored criteria beyond single-domain hits. Here, we describe the functional annotation strategies for three previously reported RIP families using hmmsearch from HMMER [36].

### Cell Division Cycle 6 Protein (CDC6)

CDC6 is the main RIP in most archaea and contains three domains: an AAA+ ATPase domain, a lid domain, and a C-terminal winged-helix (wH) domain [37]. After analyzing 797 reference sequences using InterProScan v5.77 [38], we identified the HMM profiles of the three domains: AAA+ (PF13401, PF13191), Cdc6_lid (PF22703), and WHD_Cdc6 (PF09079). Therefore, a gene was annotated as *cdc6* only if its putative protein product contained all the three domains.

### Methanococcales Putative Replication Initiation Protein (Mc-pRIP)

Mc-pRIP is a CDC6 analog in Methanococcales, as described in our previous work [39]. The helix-turn-helix (HTH_5: PF01022) and AAA+ domains of the protein were confirmed in this protein using InterProScan. However, the presence of the two domains alone is insufficient for annotation, because several ArsR family regulators also harbor the two domains [40]. In the present study, we identified a novel domain (Mc-pRIP lid) using AlphaFold 3 [41] and FoldMason [42]. Although molecular evolutionary analysis revealed a low sequence similarity between Mc- pRIP and CDC6 (Fig. 2A), their predicted structures showed high similarity, with a local distance difference test (LDDT) score of 0.546 (Fig. 2B). Furthermore, a 90-aa conserved region (Fig. 2C) between the HTH_5 and AAA+ domains matched the lid domain of CDC6 (Fig. 2D). Therefore, we inferred that this region shares a similar function with the CDC6 lid domain and built an HMM profile based on 14 reference Mc-pRIP proteins as follows. The N- and C-terminal regions were first removed using an in-house Python script. Subsequently, we performed MSA using the MUSCLE algorithm in MEGA 12 [43], trimmed the alignment using trimAI v1.5.0 [44], and constructed an HMM profile using hmmbuild from the HMMER suite. Similar to the annotation method used for *cdc6*, a gene was annotated as *mc-pRIP* if its predicted protein product contained all the three domains.

**Fig. 2.**
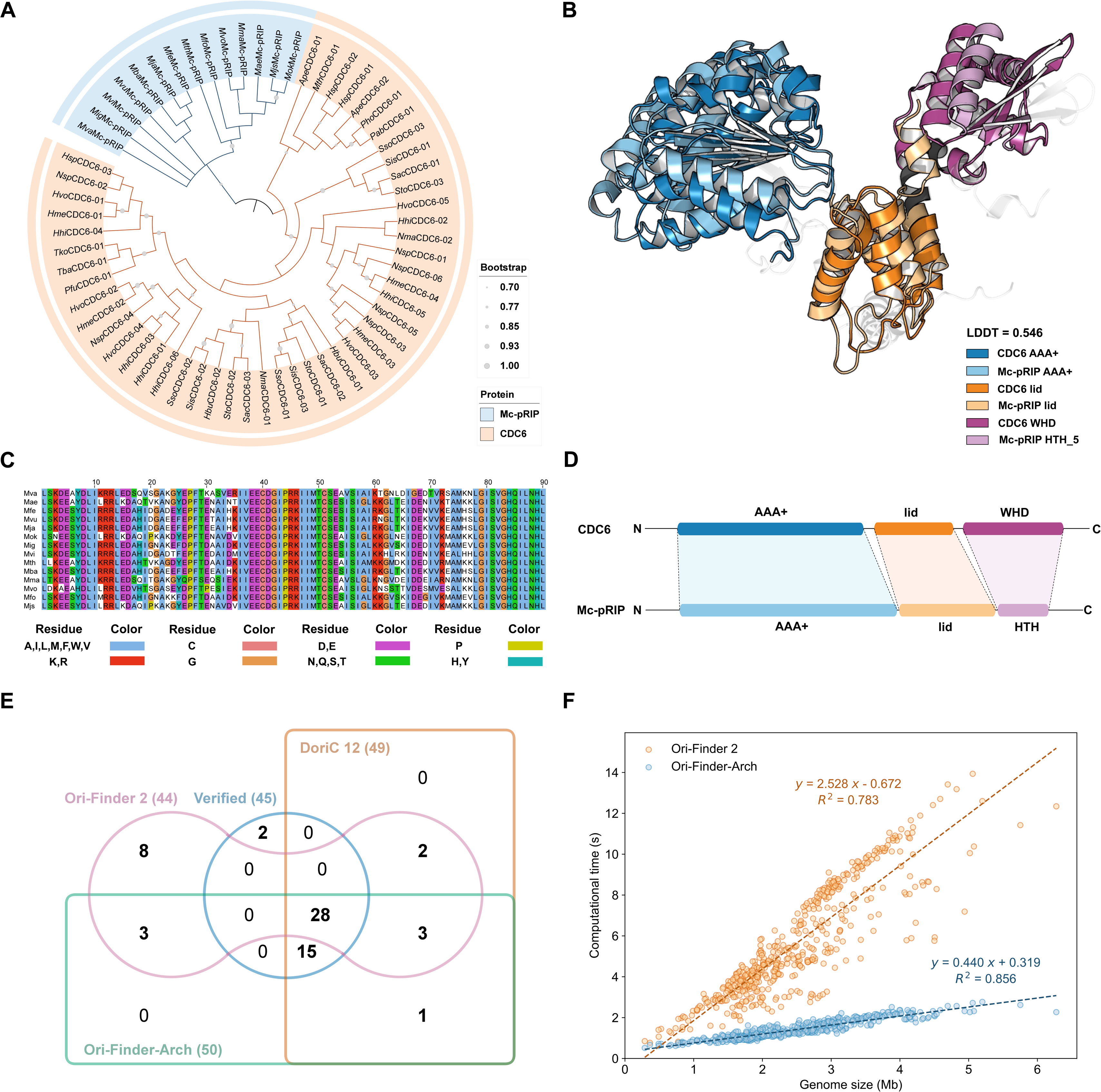
Analysis of the annotated RIPs and *oriC*s. **A.** Maximum-likelihood phylogenetic tree of representative CDC6 and Mc-pRIP. **B.** Alignment of the predicted structures of CDC6 and Mc-pRIP. It is revealed that, although Mc- pRIP and CDC6 are evolutionarily divergent, their structures are highly similar, and they are likely isofunctional. **C.** Multiple sequence alignment of the Mc-pRIP lid domains. **D.** Correspondence between the domains of CDC6 and Mc-pRIP. **E.** The relationships among the verified dataset, the DoriC dataset, and the prediction results of Ori-Finder 2 and Ori-Finder- Arch. **F.** Comparison of computational time between Ori-Finder 2 and Ori-Finder-Arch.

### Winged-helix Initiator Protein (WhiP)

WhiP is an additional RIP in *Sulfolobus* species and is considered to have two wH domains [45]. However, only a single wH domain (SSF46785) was identified in WhiP by InterProScan, and this condition is not sufficient for annotation. Therefore, we built a family HMM profile based on 53 reference WhiP proteins (the method was the same as that used for Mc-pRIP). A candidate was annotated as *whiP* if it met the criteria of a matching region ≥ 70% and an E-value ≤ 1e-3.

### Conserved Consensus Motif Recognition

#### Origin Recognition Box (ORB)

The ORB of CDC6 and Mc-pRIP consists of two distinct cis-acting elements: a dyad element recognized by the wH domain and an asymmetric G-rich motif recognized by an initiator-specific motif (ISM) within the AAA+ domain [46]. When identifying ORBs in *oriC*s from DoriC, we considered that the database might contain inaccurate predictions. Because the copy number of ORBs is often redundant, and the presence of tandem repeats (TRs) can enhance data confidence, we first screened for sequences containing TRs using Tandem Repeat Finder (TRF) [47]. Candidate ORB motifs were discovered using MEME [48], and the final motifs were determined based on visual inspection of the motif features. In our previous study, we showed that family-specific motifs outperformed common motifs in determining the *oriC*s of target strains. Therefore, in this work, we expanded the coverage of species-specific ORBs from 6 to 17 families (Table S4).

### Uncharacterized Motifs (UCM)

In the reference WhiP-type *oriC*s, we identified a previously described 12-bp inverted repeat (IR) 5′-TACTACTACTAC-3′ [45] using MEME Suite. This motif exhibits a tandem arrangement of triplets analogous in pattern to DnaA-trios (DnaA protein-binding sites in bacterial *oriC*) [49], and is used as a criterion for WhiP-type *oriC* identification. Additionally, another UCM 5′-TTATACAAAC-3′ has recently been verified in *Sulfolobus islandicus* REY15A to be associated with replication initiation [50]. Therefore, this UCM also serves as a criterion for WhiP-type *oriC* identification.

### DNA Unwinding Element Detection

A modified algorithm of GC-Profile [51] was used to identify putative DUEs. Each intergenic sequence was segmented based on local GC content variations, and the interval with a minimum length of 40 bp that exhibited the maximum slope (i.e., the most abrupt transition in GC content) was designated as the candidate DUE.

### *oriC* Annotation Pipeline

Based on these models and algorithms, we established a pipeline for archaeal *oriC* identification (Fig. 3). Firstly, Pyrodigal is used to initially annotate protein-coding genes (score ≥ 10) and IGRs (length ≥ 120 bp) in genome sequences [52]. Then, genes encoding RIPs are annotated as described above. For IGRs, ORBs and UCMs are recognized using FIMO from the MEME suite (E-value ≤ 1e-3) or regex expression (mismatch ≤ 1), and DUEs are detected using modified algorithm of GC-Profile. Note that the similarity to known *oriC*s, as determined by BLASTn (E-value ≤ 1e-10) against the experimental verified dataset or DoriC 12.0, is also serve as an optional criterion (non-mandatory) in our system. Finally, a priority decision-making system is implemented to select putative *oriC*s as follows:

**Fig. 3.**
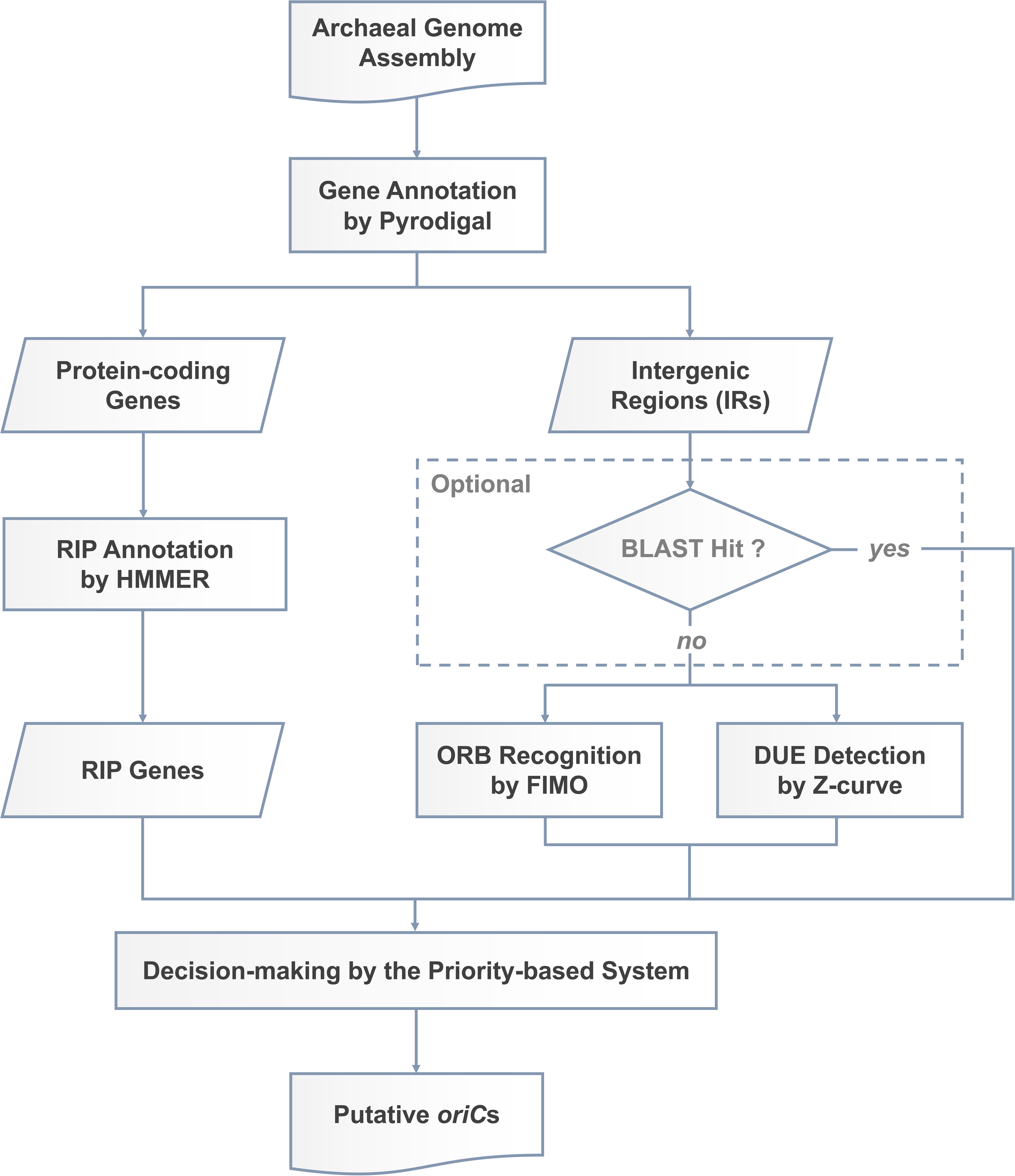
The annotation pipeline used by Ori-Finder-Arch to identify the putative *oriC*s.

**Type I: RIP-adjacent.** If an IGR lies adjacent to RIP genes and harbors ORBs or UCMs, it is designated as a putative *oriC*.

**Type II: BLAST-hit.** If BLASTn is available and no candidate satisfies the Type I criteria, any IGR that exhibits a hit to a known *oriC* in the reference database with an E-value ≤ 1e-10 and a coverage ≥ 70% is considered a putative *oriC*.

**Type III: Structure-match.** If no candidate satisfies Type I or Type II criteria, any IGR with ≥3 ORBs and the ORB(−)/DUE/ORB(+) pattern (an inverted pair of ORBs with an intervening DUE) is reported as a putative *oriC*.

These three types are prioritized in descending order (Type I > Type II > Type III) for the final *oriC* annotation.

## Results

### HMMER-based RIP annotation substantially improves *oriC* localization accuracy

On the benchmark dataset of 19 genomes with experimental evidence, we found that 70.69% of the RIPs predicted by Ori-Finder-Arch were adjacent to verified *oriC*s, compared to only 28.95% predicted by Ori-Finder 2. This indicates that the HMMER-based RIP annotation strategy is more effective for *oriC* localization than the BLAST-dependent method, which may be attributed to the ability of the domain-based method to exclude pseudogenes more effectively, allowing for the precise detection of true replication initiation-related genes. Furthermore, this method exhibited good cross-species generalizability. On the complete/chromosome-level genomes and the scaffold/contig-level genomes of the three ranks (high, medium, and low), the positive rates exhibited a downward trend, with values of 97.38%, 96.13%, 87.69%, and 68.40%, respectively, indicating that assembly continuity strongly affects RIP annotation capability.

### Ori-Finder-Arch achieves superior prediction performance to Ori-Finder 2

On the benchmark dataset of verified *oriC*s, Ori-Finder-Arch achieved a recall of 95.6% and precision of 86.0%, substantially outperforming Ori-Finder 2 (recall: 62.2%; precision: 63.6%) (Table S2). Note that BLAST was explicitly disabled for both Ori-Finder-Arch and Ori-Finder 2 in all benchmark analyses to avoid circular validation. Additionally, the speed test on complete genomes revealed that Ori-Finder-Arch ran 4.75 times faster than Ori-Finder 2 with a higher *R*²(0.856 *vs*. 0.783) (Fig. 2F; benchmarked on 32 ×Intel^®^ Core^TM^ i9-13900K, 64 GB RAM, Ubuntu 20.04), demonstrating its improved efficiency and robustness. Given its high performance, we applied Ori-Finder-Arch to all 16,409 assemblies and successfully annotated 17,472 *oriC*s across 9,877 genomes (Table S1), covering major archaeal phyla, including Methanobacteriota, Nitrososphaerota, Thermoplasmatota, Thermoproteota, Halobacteriota, Nanobdellota, and Promethearchaeota.

### The structural pattern of *oriC* is conserved in the majority of archaeal genomes

In the present study, we found that 67.0% of the predicted CDC6-type *oriC*s in the complete/ chromosome-level genomes conformed to the ORB(−)/DUE/ORB(+) pattern, and this structural pattern remained prevalent in scaffold/contig-level assemblies (58.7%) (Table S1). This observation revealed that the composition and organization of core functional elements of replication initiation are widely conserved, despite considerable variation in *oriC* numbers across different archaeal lineages. Consequently, Ori-Finder-Arch enables efficient and accurate *oriC* annotation across diverse archaeal phylogenies based on this universal principle.

### Implementation

The Ori-Finder-Arch web server is built on a modular architecture that separates the core analysis pipeline from the web interface. The backend service was developed using Flask 2.6.3 and deployed within a Docker container to ensure a consistent runtime environment. The core pipeline of Ori-Finder-Arch is written in Python and coordinates several third-party packages and external executables, including Pyrodigal 3.6.3 [52], BLAST+ 2.15.0 [19], HMMER 3.4 [36], MEME 5.5.9 [20], and Tandem Repeats Finder 4.09.1 [47]. Frontend logic was implemented in JavaScript, with sequence visualizations and annotations rendered interactively using Apache ECharts 5.5.1, Logomaker 0.8.7 [53], and Feature-Viewer 1.6.0 [54].

### Input

The Ori-Finder-Arch web server provides a newly designed data-submission interface (Fig. 4A). Users can upload genome sequence files in FASTA or GenBank format for *oriC* prediction. This pipeline accepts genome inputs at any assembly level (complete, chromosome, scaffold, or contig) and supports both circular and linear DNA sequences. Additionally, the parameters can be customized according to specific needs, including the types of RIPs (see the Methods section), BLASTn datasets, and ORB motifs (common *vs*. species-specific).

**Fig. 4.**
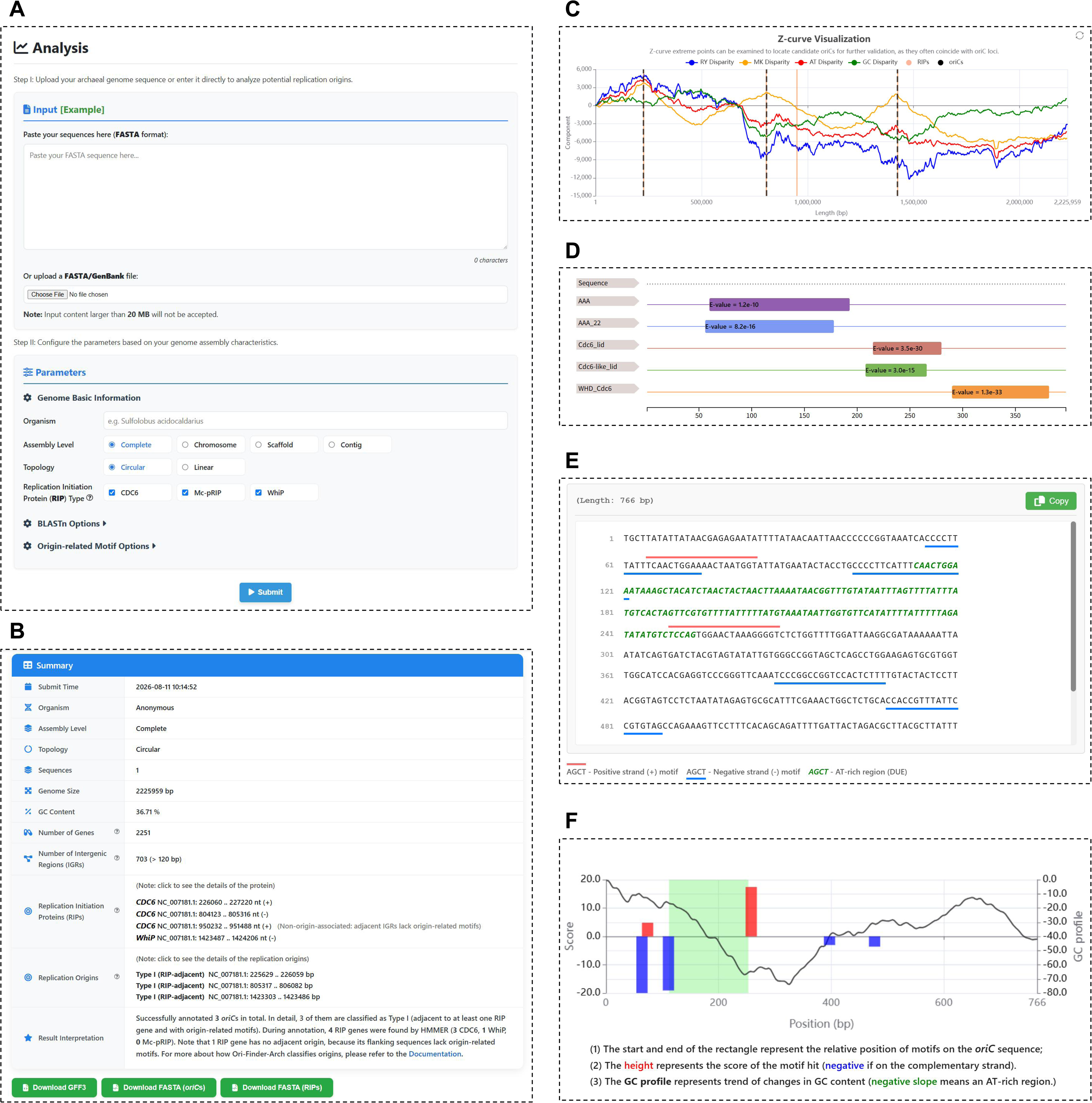
The web interface of Ori-Finder-Arch. **A.** Input submission section. **B.** Summary section of output results. **C.** Genome-wide Z-curve visualization module. **D.** Protein domain visualization module powered by Feature-Viewer. **E.** Sequence-oriented *oriC* visualization module. **F.** GC profile-based *oriC* visualization module.

### Output

The Ori-Finder-Arch web server provides both detailed text reports and interactive visualizations (Fig. 4B). At the genomic level, the relationships between *oriC* and RIP distributions and genomic composition variations are visualized using a Z-curve plot (Fig. 4C). Because the *oriC* loci in archaeal genomes tends to associate with extreme points [55], users can examine the Z-curves to identify potential *oriC*s for further confirmation.

At the element level, the relative positions and scores of the sub-elements within the proteins and replication origins are intuitively visualized. For RIPs, functional domains (e.g., AAA+ and HTH_5) are displayed as colored blocks with E-values along the amino acid sequences, with a zoom capability for close inspection (Fig. 4D). For *oriC*s, the visualization module presents each nucleotide sequence with sub-elements annotated using clear indicators: red overlines for positive-strand motifs, blue underlines for negative-strand motifs, and green italics for AT-rich regions (Fig. 4E). Additionally, a corresponding sub-element distribution plot is also provided, visualizing motif hits as rectangular blocks overlaid on a GC profile, with the height directly reflecting the FIMO score (Fig. 4F).

All the outputs can be exported and downloaded as local files in standard bioinformatics formats (GFF3 for genomic coordinates and FASTA for sequences).

## Discussion and Conclusion

Ori-Finder-Arch has significant application potential for the discovery of novel replication origins, analysis of replication initiation mechanisms, and rational design of genetic elements. Through systematic prediction across massive archaeal genomes, we identified numerous novel putative *oriC*s that substantially enrich the repository of archaeal replication origin elements. These results may include not only active *oriC*s but also potential dormant *oriC*s. For example, in *Haloferax mediterranei* ATCC 33500, Ori-Finder-Arch correctly annotated a previously verified dormant *oriC* [NC_017941.1:2069124..2069461] [56]. Furthermore, Ori-Finder-Arch not only identifies IGRs harboring *oriC*s but also provides the precise locations of functional sub-elements (ORBs, DUEs, IRs, and UCMs). This capability facilitates the theoretical design of minimal replication origin elements conforming to either the ORB(−)/DUE/ORB(+) pattern or the IR(+)/UCM/IR(−) pattern (WhiP-type). After rigorous curation, these data can serve as high-quality training sets for generative models (e.g., OriGen [57]), thereby guiding the rational design of synthetic replication origin elements.

Nevertheless, Ori-Finder-Arch still has some limitations. First, its performance is constrained by the accuracy of gene annotation. For example, in *Saccharolobus islandicus* M.16.4, Pyrodigal predicted a putative gene with a low score [NC_012726.1:486..767(+)], which overlaps with an experimentally verified *oriC* [50], thereby interfering with the correct *oriC* annotation. Future improvements could involve integrating more accurate gene annotation algorithms (e.g., GeneMarkS-2+ [58]) to reduce the impact of false-positive gene calls. Second, application of Ori-Finder-Arch to draft genomes and metagenomes remains challenging. As demonstrated in the Results section, the recall rate decreased significantly for highly fragmented contigs. Future work could leverage long-context large language models (e.g., Evo [59]) to infer contig adjacency relationships and repair low-continuity sequencing regions, thereby further enhancing *oriC* identification performance in complex genome assemblies.

In conclusion, Ori-Finder-Arch provides an efficient, accurate, and user-friendly platform for the scientific community to study archaeal replication initiation. We believe that the Ori-Finder-Arch web server will be a valuable resource for analyzing the mechanisms of DNA replication initiation in archaea, driving new discoveries and advances in biological applications. In addition, this novel architecture offers valuable guidance for performance optimization of the entire Ori-Finder system. Therefore, we plan to generalize this framework to other Ori-Finder modules, such as Ori-Finder 2022 [62] and OriV-Finder [63], and develop a unified access interface in the future. Such an extension would not only broaden its applicability, but also greatly improve its accessibility and practical value for the broader research community.

## Supporting information

Prioritized selection algorithm of archaeal oriCs

Statistics of archaeal genomes and predicted oriCs

Comparison of prediction performance on the validated dataset

Reference replication initiation proteins with InterPro domain annotations

Common and family-specific ORB motifs for archaeal oriC prediction

## Data availability

The genome sequences and annotation data were directly from the NCBI and DoriC databases and can be accessed via the accession numbers provided in Table S2. Other data are incorporated into the article and its supplementary material.

## Code availability

The web server, together with stand-alone software, model files, and prediction result files (GFF3 format), is freely available at https://tubic.org/Ori-Finder-Arch/ and https://tubic.tju.edu.cn/Ori-Finder-Arch/. This software can also be downloaded from BioCode (Accession: BT008164).

## CRediT author statement

**Zhisong You:** Data curation, Statistical analysis, Methodology, Validation, Writing-original draft, Visualization. **Zetong Zhang:** Software, Statistical analysis, Methodology, Validation, Writing-original draft, Visualization. **Hao Luo:** Conceptualization, Writing-review & editing, Supervision, Funding acquisition. **Feng Gao:** Conceptualization, Writing-review & editing, Supervision, Funding acquisition. All the authors have read and approved the final manuscript.

## Acknowledgements

We thank Yujie Li for technical assistance and beneficial discussions throughout this work. This work was supported by the National Natural Science Foundation of China (grant numbers 32270692, 31801104, and 31571358) and the Independent Research Project of State Key Laboratory of Synthetic Biology (Project No. HCZB-202601A).

## Supplementary Material

**File S1 Prioritized selection algorithm of archaeal *oriC*s**

**Table S1 Statistics of archaeal genomes and predicted *oriC*s**

**Table S2 Comparison of prediction performance on the validated dataset**

**Table S3 Reference replication initiation proteins with InterPro domain annotations**

**Table S4 Common and family-specific ORB motifs for archaeal *oriC* prediction**

