## Supplementary material for "Ori-Finder-Arch: An Updated Web Server for the Annotation and Visualization of Archaeal Replication Origins": Prioritized selection algorithm of archaeal oriCs

**File S1**

| **Algorithm :** | | Select replication origins from candidate intergenic regions |
| --- | --- | --- |
| **Input** **:**  **Output :** | | Intergenic regions with RIP adjacency, ORB hits, BLASTn hits and AT-rich region information.  Chromosomal replication origins (oriCs). |
| 1 | // Determine whether an intergenic region contains the Type III oriC | |
| 2 | FUNCTION is_type_iii(*intergenic*) | |
| 3 | IF *intergenic.at_rich* = NULL OR *intergenic.motif_hits.size* < 3 | |
| 4 | RETURN FALSE | |
| 5 | END IF | |
| 6 | *at_rich* ← *intergenic.at_rich* | |
| 7 | *pos_hits* ← FILTER *intergenic.motif_hits* WHERE *strand* = +1 | |
| 8 | *neg_hits* ← FILTER *intergenic.motif_hits* WHERE *strand* = -1 | |
| 9 | IF *pos_hits.size* = 0 OR *neg_hits.size* = 0 | |
| 10 | RETURN FALSE | |
| 11 | END IF | |
| 12 | *left* ← MIN { *hit.start* FOR *hit* IN *neg_hits* } | |
| 13 | *right* ← MAX { *hit.end* FOR *hit* IN *pos_hits* } | |
| 14 | IF *left* < *right* AND *right* > *at_rich.start* AND *left* > *at_rich.start* // Intersection | |
| 15 | *overlap* ← MIN { *at_rich.end, right* } *-* MAX { *at_rich.start, left* } | |
| 16 | *atr_ratio* ←  *overlap* / *at_rich.length* | |
| 17 | *orb_ratio* ←  *overlap* / (*right - left*) | |
| 18 | IF *atr_ratio* > 0.8 AND *orb_ratio* > 0.2 | |
| 19 | RETURN TRUE | |
| 20 | END IF | |
| 21 | END IF | |
| 22 | RETURN FALSE | |
| 23 | END FUNCTION | |
| 24 | // Select oriCs by the priority-based system | |
| 25 | FOR *intergenic* IN *intergenics* DO: | |
| 26 | // Type I: RIP-adjacent oriC | |
| 27 | IF *intergenic.adjacent_to_rip* AND *intergenic.motif_hits.size* > 0: | |
| 28 | YIELD *intergenic* | |
| 29 | // Type II: BLAST-Hit oriC | |
| 30 | ELSE IF *intergenic.blastn_hits.size* > 0 | |
| 31 | YIELD *intergenic* | |
| 32 | // Type III: Structually matched oriC | |
| 33 | ELSE IF is_type_iii(*intergenic*) | |
| 34 | YIELD *intergenic* | |
| 35 | END IF | |
| 36 | END FOR | |
