## Supplementary material for "Ori-Finder-Arch: An Updated Web Server for the Annotation and Visualization of Archaeal Replication Origins": Common and family-specific ORB motifs for archaeal oriC prediction

**Table S4**

| **Motif_ID** | **Target_Group** | **Motif_Sequence** | **E-value** |
| --- | --- | --- | --- |
| Common_CDC6 | CDC6 | 5’-TTACACYGGAAACGN**GGGG**KK-3’ | 1.7E-607 |
| Common_WhiP | WhiP | 5’-TACTACTACTAC-3’ | 2.0E-116 |
| Common_Mc-pRIP | Mc-pRIP | 5’-TTACACYGGAAACGN**GGGG**KK-3’ | 5.9E-023 |
| Family01_Haloferacaceae | Haloferacaceae | 5’-TTTCACCGGAAACGA**GGKGTGTG**TS-3’ | 3.6E-692 |
| Family02_Natrialbaceae | Natrialbaceae | 5’-TTWCACCGGAAACGC**GGKGTGTG**-3’ | 6.7E-107 |
| Family03_Sulfolobaceae | Sulfolobaceae | 5’-CGGTATTTYTCCMGTGGAAATARA**GGGG**-3’ | 1.7E-084 |
| Family04_Haloarculaceae | Haloarculaceae | 5’-AMMACTGRMACGGT**GGKGTGTG**-3’ | 6.0E-236 |
| Family05_Methanosarcinaceae | Methanosarcinaceae | 5’-NATACAGTNGAARCAAA**GGGGG**CAAAGCGT-3’ | 1.5E-272 |
| Family06_Nitrososphaeraceae | Nitrososphaeraceae | 5’-WTWTAWAGTTCCANTGGAAATAAA**GGGG**T-3’ | 6.8E-046 |
| Family07_Halobacteriaceae | Halobacteriaceae | 5’-CNGNWACASYGGAAACRGT**GGKGTGGGGG**-3’ | 1.8E-048 |
| Family08_Nitrosopumilaceae | Nitrosopumilaceae | 5’-CMTGGAGTGGAAATAAA**GGKG**TGTGAGT-3’ | 1.5E-276 |
| Family09_Haladaptataceae | Haladaptataceae | 5’-CNNRWTACASCGGAAAC**GGGGGGG**T-3’ | 2.4E-047 |
| Family10_Methanoregulaceae | Methanoregulaceae | 5’-TCCACTGGAAACAAA**GGG**-3’ | 7.2E-153 |
| Family11_Methanotrichaceae | Methanotrichaceae | 5’-TCCAGTGGAAAYMAW**GGGG**-3’ | 2.7E-128 |
| Family12_Methanomethylophilaceae | Methanomethylophilaceae | 5’-TTCCASTGGAAATRGA**GGGG**T-3’ | 3.0E-057 |
| Family13_Natronomonadaceae | Natronomonadaceae | 5’-GTTCCACCCGAAACGAR**GGGGTGG**-3’ | 2.7E-049 |
| Family14_Methanomicrobiaceae | Methanomicrobiaceae | 5’-NAGAGWTCCAGWGGAAACAAA**GGGG**TCRGG-3’ | 5.3E-055 |
| Family15_Methanocorpusculaceae | Methanocorpusculaceae | 5’-RRKAACTGCAGAACARC**GGCNGGG**-3’ | 2.6E-028 |
| Family16_Halorubellaceae | Halorubellaceae | 5’-GTTYCACTYGAAAC**GGTGGKGKGGGGG**NN-3’ | 4.5E-021 |
| Family17_Natronoarchaeaceae | Natronoarchaeaceae | 5’-AACACYGGAAAC**GGTGGGGTGG**-3’ | 4.1E-028 |

Note: This table presents a manually curated set of ORB motifs derived from DoriC entries. To enhance data confidence, we first screened for tandem repeats (TRs) using Tandem Repeat Finder (TRF). Candidate motifs were then discovered with MEME and finalized through visual inspection of motif features. The motifs are organized into three common types (CDC6, WhiP, Mc-pRIP) and 17 family-specific groups, expanding our previous coverage from 6 to 17 archaeal families. These motifs can be used to improve the accuracy of replication origin prediction in target strains.
